# Structural bioinformatics of three Epstein-Barr Virus (EBV) Integral Membrane Proteins and their water-soluble QTY analogs

**DOI:** 10.64898/2026.07.22.740197

**Authors:** Zixuan Sun, Edward Chen, Shuguang Zhang

## Abstract

The Epstein-Barr virus (EBV) is a highly prevalent virus worldwide that is associated with several lymphoid and epithelial malignancies. However, extensive research on EBV integral membrane proteins BILF1, LMP1 and LMP2, has been scarce due to their hydrophobic transmembrane domains. Our study applies the QTY code (glutamine, threonine, tyrosine) to design water-soluble analogs of BILF1, LMP1 and LMP2 with reduced hydrophobicity, where we systematically replaced hydrophobic amino acid residues leucine (L), isoleucine (I), valine (V), and phenylalanine (F) with structurally similar polar residues glutamine (Q), threonine (T), and tyrosine (Y). We retrieved their native sequences from UniProt, identified transmembrane domains using Protter, then performed QTY design through the Protein Solubilizing Server (PSS). We then predicted native and QTY structures using *in silico* prediction tools AlphaFold3, ColabFold, and Boltz-2. Our analyses demonstrate that despite significant protein sequence replacements in their transmembrane domains (54.15%-61.59%) and increased intrinsic solubility, the QTY analogs exhibited minimal changes in isoelectric point (0.00-0.15 decrease) and molecular weight (0.7-1.2 kDa increase). Additionally, structural superpositions between QTY analogs and native structures using PyMOL yield low RMSD values (0.217Å −1.202Å). Our results demonstrate the QTY code’s ability to design detergent-free analogs of BILF1, LMP1 and LMP2 with substantially reduced hydrophobicity and aggregation propensity whilst preserving native-like structures. Our results may facilitate protein characterization studies, therapeutic research on EBV, and other protocols that typically require protein solubilization.

## Introduction

The Epstein-Barr virus (EBV), also known as human herpesvirus 4 (HHV4), is a ubiquitous, lymphotropic γ-herpesvirus infecting 90-95% of the global adult population, establishing lifelong latency primarily in B lymphocytes [1, 2]. EBV is classified as a Group 1 carcinogen by the World Health Organization due to its strong association with the pathogenesis of various severe lymphoid and epithelial cancers [2]. EBV’s association with human cancers was first established as early as 1964, as EBV was the first virus isolated from a human tumor – specifically, Burkitt’s lymphoma [2]. Furthermore, EBV is evidently linked to infectious mononucleosis, Hodgkin’s lymphoma, nasopharyngeal carcinoma (NPC), post-transplant lymphoproliferative disease (PTLD), and gastric carcinoma [3]. Globally, EBV-associated malignancies account for approximately 1.9% of all human cancers, thus resulting in around 357,900 new cancer cases annually [4]. However, despite EBV’s early discovery and the significant threat it poses to public health worldwide, therapeutic and drug research on EBV remains challenging [5].

Our current structural bioinformatic study focuses on three EBV integral membrane proteins that are involved in its oncogenesis, immune evasion and latency: the viral G protein-coupled receptor (GPCR) BILF1 [1, 6], Latent Membrane Protein 1 (LMP1), Latent Membrane Protein 2 (LMP2).

BILF1 (UniProt ID: P0DXO9) is a GPCR with seven alpha-helical transmembrane domains that plays a key role in EBV immune evasion, possessing the notable ability of disrupting the cell surface expression of major histocompatibility complex class I (MHC-I) molecules [6, 7]. Specifically, BILF1 physically associates with MHC-I molecules on the cell surface and promotes their internalization through its EKT motif near transmembrane helix 3. Consequently, BILF1’s cytoplasmic C-terminal tail contributes to the degradation of internalized MHC-I molecules by promoting the lysosomal pathway [8, 9]. Additionally, a 2011 study found that BILF1 interferes with the exocytic pathway, thereby inhibiting the localization of MHC-I and further decreasing its cell-surface concentration. Since cell-surface MHC-I molecules are directly involved in the detection of EBV-infected cells by CD8^+^ cytotoxic T cells, their removal by BILF1 allows for immune evasion[6, 8]. Moreover, as a viral GPCR with constitutive G_i_ signaling activity, BILF1 also shows capabilities in activating oncogenesis-associated pathways in host cells [7, 10].

LMP1 (UniProt ID: P03230) is considered the primary EBV oncoprotein, being a 66kDa integral membrane protein composed of a short 24-amino acid cytoplasmic N-terminus, six transmembrane (TM) domains, and a 200-amino acid cytoplasmic C-terminal signaling tail [11]. Functionally, LMP1 acts as a constitutively active analog of the human costimulatory CD40 receptor, a member of the Tumor Necrosis Factor Receptor (TNF-R) family [12, 13]. The CD40 receptor is primarily expressed on various antigen-presenting cells, including B cells, dendritic cells, and macrophages [14]. In B cells, the primary targets of EBV, CD40 signaling is closely associated with cell survival and proliferation in adaptive immune responses. During these responses, receptors on antigen-specific B cells first bind with extracellular antigens, then internalize and process them, and ultimately display resulting peptide fragments on major histocompatibility complex (MHC) class II molecules. These peptide-MHC II complexes are recognized by activated CD4^+^ helper T cells, which express CD40L on their cell membranes. Consequently, CD40L binds to CD40, activating multiple downstream pathways in the B cell, most notably NF-κB and MAPK/JNK [14]. These pathways promote B cell survival, germinal center formation, proliferation, and eventual differentiation into memory B cells and plasma cells [15]. Therefore, by mimicking CD40 signaling, LMP1 subverts apoptotic pathways whilst promoting proliferation of EBV-infected B cells, contributing to EBV-associated oncogenesis through B-cell transformation [11, 12].

LMP2 (UniProt ID: P13285) is less characterized than LMP1 and BILF1. It lacks a complete Cryo-EM structure on both UniProt and the RCSB Protein Data Bank (PDB) as of June, 2026. Current research confirms that LMP2 exists mainly as two isoforms, LMP2A and LMP2B [16]. These isoforms have identical transmembrane regions consisting of twelve α-helices, and their major difference comes from LMP2A having an additional 119-amino-acid cytoplasmic signaling domain at its N-terminus that is absent in LMP2B [11]. Since our protein designs and analyses pertain only to the shared TM regions of the isoforms, subsequent analyses will use “LMP2” to refer to this common transmembrane region. Functionally, LMP2A is regarded as the main signaling isoform, partially mimicking B-cell receptor (BCR) signaling whilst suppressing normal BCR activation in EBV-infected B cells. These functions allow LMP2A to activate downstream pathways associated with cell survival and proliferation, such as the PI3K/Akt pathway, thereby supporting EBV latency and host-cell survival without triggering the complete BCR response that could lead to EBV lytic reactivation [16, 17, 18].

LMP2B is less characterized than its other isoform, potentially due to its lower expression than LMP2A in certain malignancies such as NPC [11]. Regardless, it is currently believed that the isoform could inhibit LMP2A’s N-terminal signaling activities by forming heterodimers with it. This inhibition may counteract LMP2A’s role in maintaining EBV latency, thereby promoting lytic replication [19, 20].

BILF1, LMP1, and LMP2 are all EBV integral membrane proteins with varying numbers of alpha-helical TM domains. The inherent hydrophobicity of TM domains is a significant barrier to characterizing these proteins, as they typically need to be solubilized by detergents before extraction or crystallization [21]. Therefore, given the high costs and long development times of detergents, this study aims to design water-soluble analogs of BILF1, LMP1, and LMP2, using structural bioinformatics approaches [22].

Instead of replacing amino acids arbitrarily, we apply the QTY code, a very specific amino acid replacement method for designing water-soluble analogs of transmembrane proteins with reduced hydrophobicity via systematic amino acid substitution. The rationale for these replacements is the high structural similarity between these hydrophobic and polar amino acid pairs observed in high-resolution electron density maps [23]. Specifically, the QTY code replaces leucine (L) with glutamine (Q), isoleucine (I)/valine (V) with threonine (T), and phenylalanine (F) with tyrosine (Y). Utilizing the QTY code to solubilize transmembrane proteins has been successful in numerous prior studies, including Zhang *et al.*’s initial application to design four detergent-free chemokine receptors, all of which retained ligand-binding activities, thermostabilities, and tertiary structures despite significant changes in hydrophobicity [23–25]. Other studies include *in silico* analyses of QTY analogs of glutamate transporters, ABC transporters, solute carrier transporters, integral membrane enzymes, NADPH oxidases, and monoamine neurotransmitter transporters [26–31]. Furthermore, past studies have designed biomimetic sensing systems and molecular traps targeting cancer metastasis using similar methods [32, 33]. These studies demonstrate the broad applicability of the QTY code.

Our current study applies the QTY code to BILF1, LMP1, and LMP2. We did protein sequence alignments, used AlphaFold 3 for structure prediction, then and water solubility analysis to evaluate whether QTY analogs of these EBV integral membrane proteins preserve their native structures while becoming water-soluble.

## Results and Discussion

### Sequence alignments and comparisons of other characteristics of EBV integral membrane proteins

The amino acid sequences of three EBV integral membrane proteins were aligned with sequences of their QTY analogs (Figure 1). After application of the QTY code, variations in overall sequences ranged from 22.02% to 27.88%, while variations in the transmembrane domains (TMD) ranged from 54.15% to 61.59% (Table 1). The considerably greater variations in TMD residues are consistent with the rationale of the QTY code – solubilizing a protein through systematic replacing hydrophobic residues LIVF in transmembrane domains with hydrophilic residues QTY [23].

**Figure 1.**
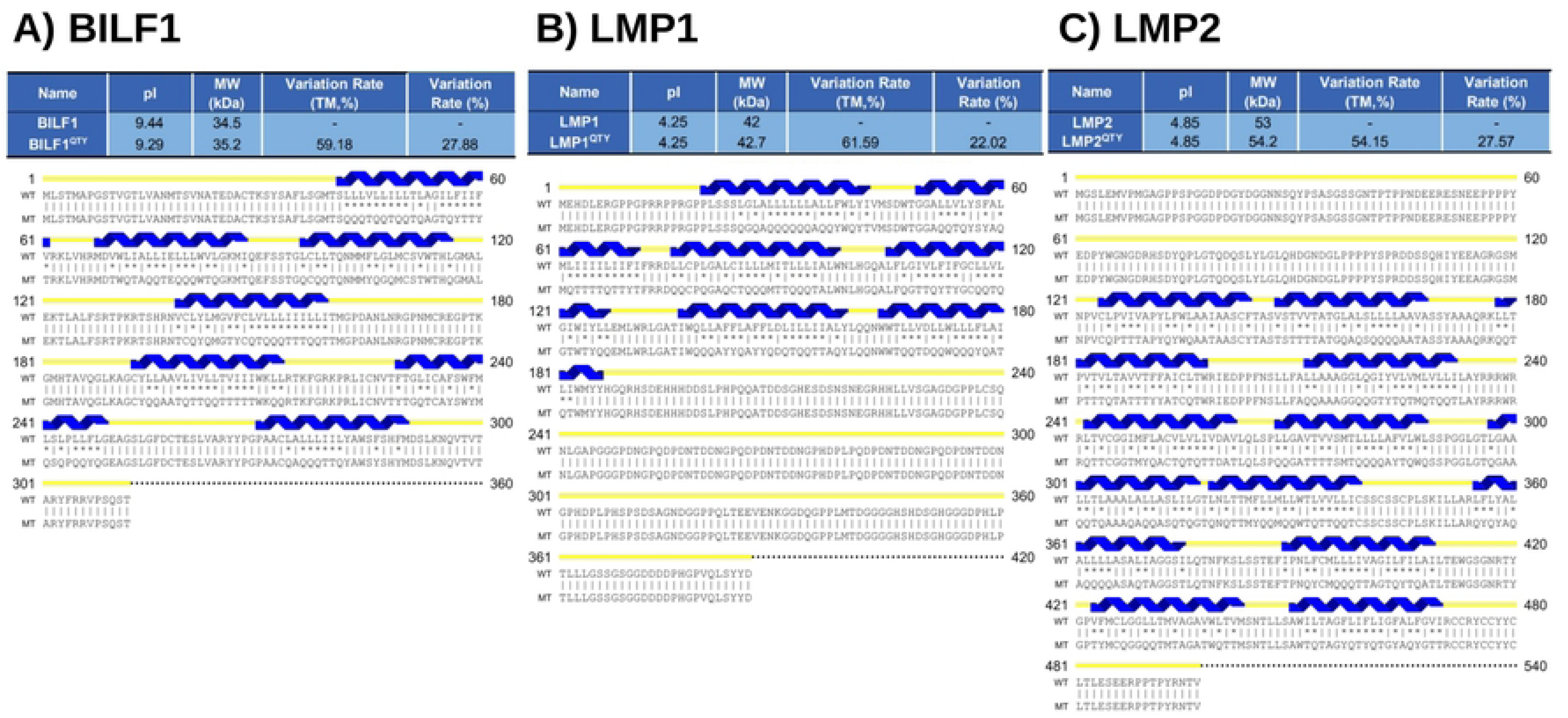
Protein sequence alignments of three native (*Upper*) EBV integral membrane proteins with their QTY analogs (*Lower*). The symbols | and * indicate whether amino acids are identical or different, respectively. Transmembrane (TM) alpha helices are shown above the protein sequences (colored in blue). The QTY code replaces the hydrophobic residues L, I/V, and F in TM helices with hydrophilic residues Q, T, and Y, respectively. Additionally, the isoelectric points (pI), molecular weights (MW), transmembrane variation %, and total variation % of the native and QTY variants are listed above the protein sequences. The alignments are A) BILF1 versus BILF1^QTY^, B) LMP1 versus LMP1^QTY^, C) LMP2 versus LMP2^QTY^. Despite significant sequence variations in their TM regions (ranging from 54.15% to 61.59%), the QTY variants mostly preserve the pI and MW of the native proteins.

**Table 1.**
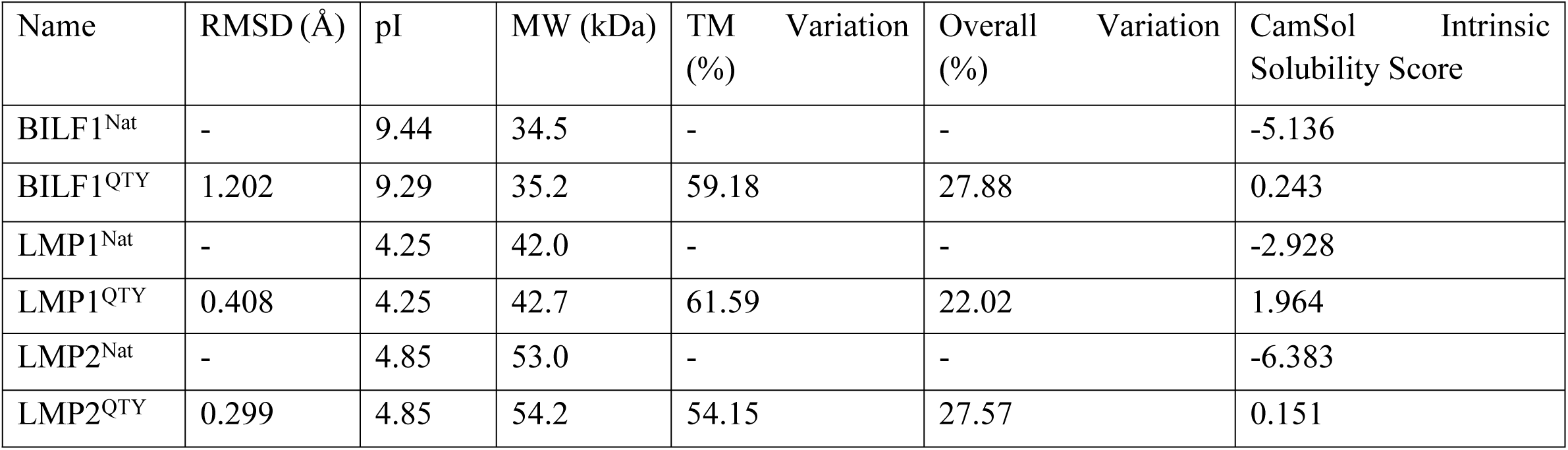
Protein characteristics of three Epstein-Barr virus integral membrane proteins and their QTY analogs.

Despite significant replacements in amino acid sequences, the proteins’ isoelectric points (pI) experienced minimal decreases, with changes ranging from 0.00 to 0.15 (Table 1). This is because all amino acids involved in the QTY code, glutamine (Q), threonine (T), tyrosine (Y), isoleucine (I), leucine (L), and phenylalanine (F), are electrically neutral.

Additionally, the increases in molecular weight (MW) were minimal, ranging from 0.7kDa to 1.2kDa (Table 1). These negligible variations in overall MW are due to the similar masses of substituted residues, specifically: L (131.17 Da) and Q (146.14 Da); I (131.17 Da), V (117.15 Da) and T (119.12 Da); F (165.19 Da) and Y (181.19 Da).

### AlphaFold 3 structural predictions

AlphaFold 3 (https://alphafoldserver.com/), developed and released by Google DeepMind and Isomorphic Labs in May 2024, is among the most advanced computer tools for biomolecular structure prediction, capable of modeling proteins, nucleic acids, ligands, ions, and modified residues [34]. Following the breakthrough performance of AlphaFold 2 at CASP14, AlphaFold 3 performed strongly in CASP16, particularly in tasks involving protein and other biomolecular complexes [35, 36].

In our current study, AlphaFold 3 was used as the primary structural prediction tool, while ColabFold and Boltz-2 were employed when AlphaFold 3 failed to generate reliable models. Particularly, AlphaFold 3 failed to consistently generate models that fall within the confident range of 70 < pLDDT < 90 for LMP1, LMP2, and their corresponding QTY analogs. This difficulty highlights the current challenges in modeling viral transmembrane proteins, which likely stem from the limited number of experimentally determined structural analogs and the flexibility or disorder at their termini. These limitations suggest that further improvements in computational models are still needed for accurately modeling highly modified viral membrane proteins.

### ColabFold

ColabFold(https://colab.research.google.com/github/sokrypton/ColabFold/blob/main/AlphaFold2.ip <u>ynb</u>), was released by Mirdita *et al*. in 2022 and hosted on Google Colaboratory in 2022, is an optimized implementation of AlphaFold 2 and RoseTTAFold that can greatly reduce prediction times and computational costs by replacing the slow, conventional step of multiple sequence alignment (MSA) search with the more efficient MMSeqs2 algorithm [37]. In this study, ColabFold was used to perform template-based modeling for proteins for which AlphaFold3 failed to produce reliable models.

The high pTM, ipTM, and pLDDT scores of models obtained via template-based modeling indicate that the engineered QTY analogs can adopt tertiary structures that closely resemble their native counterparts [36]. This indicates that the QTY substitutions do not introduce severe steric clashes or major structural disruptions to the native fold. However, because these results were obtained through template-based modeling, we recognize that they do not definitively establish that the QTY analogs will autonomously fold into these conformations, nor do they prove that these are the most energetically favorable states of the QTY analogs.

### Boltz-2

Boltz-2 (https://build.nvidia.com/mit/boltz2) was released by Passaro *et al.* in June 2025, is a computational biomolecular prediction model designed for both structure and binding-affinity predictions [38]. In this study, Boltz-2 was used as an alternative prediction tool when AlphaFold 3 failed to produce a reliable structure of the native LMP1 monomer, thereby enhancing the overall accuracy of the resulting protein models.

AI tools have recently been penetrating bioinformatic widely, despite using multiple computational tools to maximize the accuracy of predicted protein models, experimental validation remains crucial for confirming our computational predictions.

### Accuracy assessment of in silico predictions

The accuracy of the predicted protein structures was assessed using multiple confidence metrics generated by AlphaFold 3, ColabFold, and Boltz-2. Metrics include the per-residue scores of the predicted Local Distance Difference Test (pLDDT), predicted aligned error (PAE), and predicted Template Modeling score (pTM). Additionally, the interface predicted Template Modeling score (ipTM) was evaluated for the predicted multimers (Figure 2).

**Figure 2.**
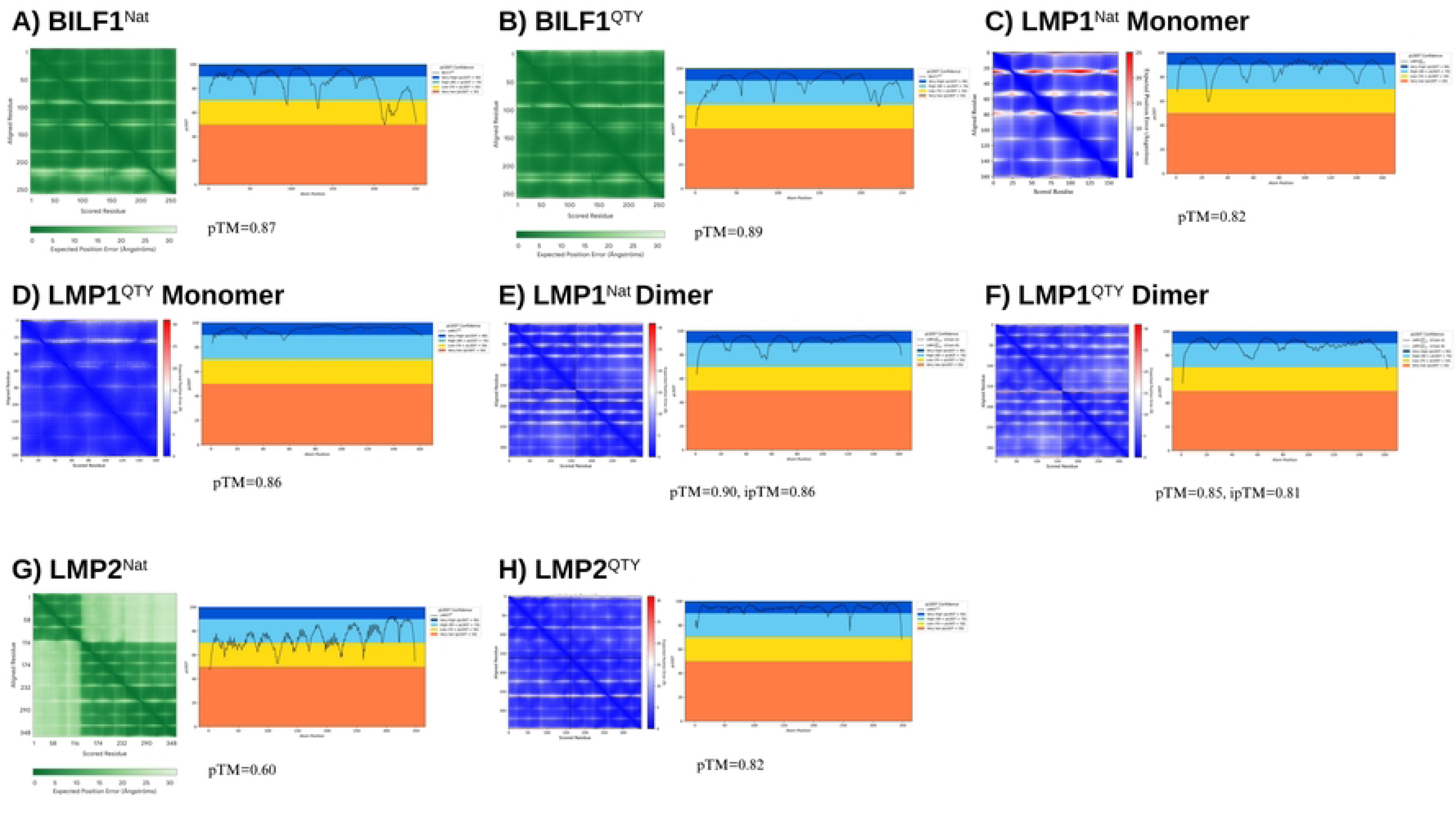
*In silico* confidence metrics (pLDDT, PAE, pTM) for eight predicted native and QTY-analog EBV integral membrane protein models. pLDDT confidence profiles, PAE matrices, and pTM scores are displayed for each predicted structure to validate topological accuracy. To ensure predictive accuracy, different AI-based structural prediction models were used. **(A-B, G)** Native and QTY variants of BILF1, alongside native LMP2, were predicted using AlphaFold 3. **(C)** The native LMP1 monomer was predicted using Boltz-2. **(D, E-F, H)** The QTY LMP1 monomer, both LMP1 homodimers, and the QTY LMP2 variant were predicted via template-based modeling in ColabFold v1.6.1. Scores of the predicted Local Distance Difference Test (pLDDT) suggest TM domains were predicted with predominantly high to very high confidence (blue and dark blue shaded areas), with limited regions showing lower confidence (*yellow and red*). Predicted aligned error (PAE) matrices show predominantly high confidence (*dark green or dark blue*) in the global 3D structures. However, the native LMP2 model shows extended regions of low confidence in both plots, indicating greater structural uncertainty. Overall, high predicted Template Modeling (pTM) and interface pTM (ipTM) scores indicate high prediction accuracies of the global structure and the docking interface between dimer chains, respectively. A) BILF1^Nat^, B) BILF1^QTY^, C) LMP1^Nat^ Monomer, D) LMP1^QTY^ Monomer, E) LMP1^Nat^ Dimer, F) LMP1^QTY^ Dimer, G) LMP2^Nat^, H) LMP2^QTY^

The pLDDT score reflects the computational model’s confidence in the predicted position of each residue. In the pLDDT confidence profiles (Figure 2), dark blue areas (pLDDT > 90) indicate very high confidence; blue areas (90 > pLDDT > 70) represent confident regions; yellow areas (70 > pLDDT > 50) signify low confidence; and orange areas (pLDDT < 50) show very low confidence. These pLDDT ranges and their corresponding confidence levels are consistent with those displayed on the AlphaFold3 web server [36]. In this study, the flexible or disordered C- and N-terminal regions generally exhibited low pLDDT scores and significantly lowered pTM values of the predicted models. These low-confidence regions were therefore truncated for clarity before structural predictions to reduce noise in RMSD analysis and better reflect the prediction accuracy for transmembrane domains. This decision is also justified by the fact that terminal regions of integral membrane proteins naturally exist in aqueous environments and therefore are not targeted by the QTY code. It is very common that during experiments, many unstructured protein termini were removed before protein synthesis and expressions.

PAE was further used to assess confidence in the relative positioning of residue pairs. It is represented as a matrix in which each point corresponds to the predicted positional error between two residues [36]. Low PAE values, represented in dark green or blue, indicate higher confidence, whereas high PAE values, represented in light green or red, indicate greater uncertainty. Our PAE matrices generally showed low predicted errors, with higher PAE values primarily associated with flexible inter-helical loop regions.

The pTM was evaluated to assess overall confidence in the predicted model, with values ranging from 0 to 1, where higher values indicate better structural alignment between the predicted model and the true structure and, therefore, greater confidence in the global protein structure. For multimeric models, ipTM was additionally used to assess confidence in predicted interchain interfaces.

Overall, most models obtained in this study exhibited high pLDDT, low PAE, high pTM, and, when applicable, high ipTM values (Figure 2), indicating high regional and global confidence in the predicted models, thereby confirming their reliability for further analysis. The alpha-helical TM domains generally exhibited higher confidence than flexible helical-loop regions, a normal phenomenon that will be further elaborated in the following paragraphs.

However, the predicted models for native LMP2 and native BILF1 require closer interpretation as reflected in one or more metrics. First, the native LMP2 model contains numerous short, low-pLDDT regions in its pLDDT confidence profile (Figure 2). These regions appear to mostly correspond to the flexible inter-helical loops connecting TM helices predicted by Protter, including residues S20–V29, Y49–T58, R77–L87, A112–T121, D138–P145, S167–L175, L194–T200, S218–R232, A249–N269, T289–R296, and T319–S327 in the truncated sequence. Hence, the prediction accuracy of its TM domains is likely higher than that reflected in the pLDDT confidence profile of the entire truncated protein. However, the PAE matrix of this model contained relatively large light-green regions, indicating moderate uncertainty in the relative positioning of multiple residue pairs. This is consistent with its relatively low pTM score of 0.60. Therefore, results derived from the predicted native LMP2 model should be interpreted with greater caution.

Additionally, in the native BILF1 model, a section of residues corresponding to TM domains 6 and 7 also exhibited low pLDDT scores. However, the PAE matrix of this model was predominantly dark green, suggesting high confidence in the relative positioning of residue pairs across the model. This is further supported by its high pTM score of 0.87. Therefore, despite the localized uncertainty in the TM6-TM7 region, there is strong confidence in the overall predicted structure of native BILF1.

### Superposed structures of native EBV integral membrane proteins and their computationally predicted water-soluble QTY analogs

We superposed the computationally predicted native structures of the EBV integral membrane proteins, along with the available experimentally determined Cryo-EM models, with their respective QTY analogs (Figure 3). Additionally, since LMP1 tends to assemble into homodimers, we also predicted and superposed structures for LMP1 dimers [12]. These superpositions enable direct comparisons between the structures of native proteins and their water-soluble QTY analogs.

**Figure 3.**
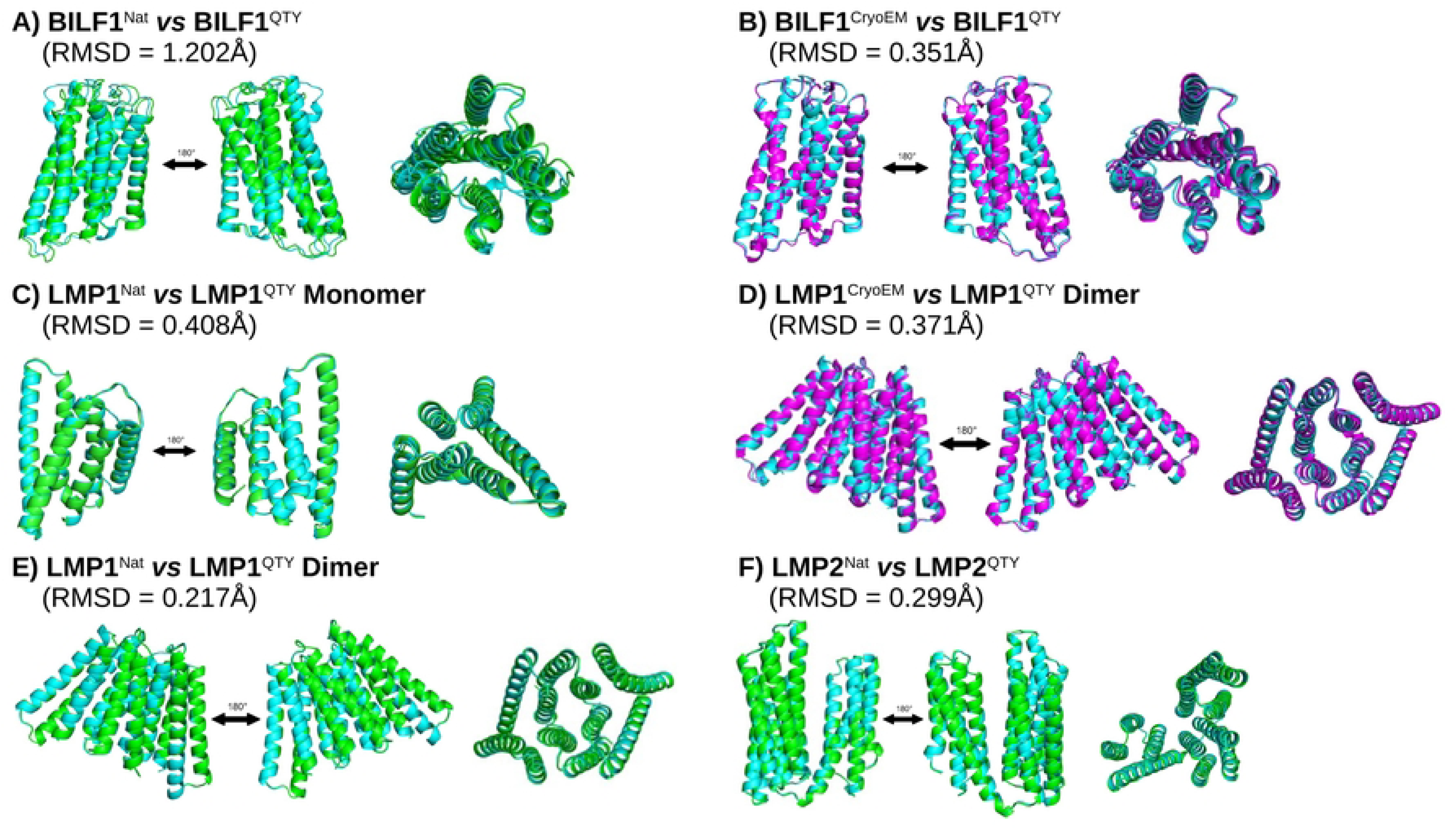
Pairwise structural superimpositions of native, QTY-analogs, and Cryo-EM models for three EBV integral membrane proteins. Computationally predicted QTY analogues (*cyan*) of three membrane proteins were compared to their corresponding predicted native structures (*green*). Comparisons were also made with Cryo-EM-determined native structures (*magenta*), when available. All Cryo-EM structures were obtained from the Protein Data Bank (PDB). Protein sequences coding for the disordered C- and N-termini were removed before computational predictions to improve the clarity of the models. The root-mean-square distance (RMSD) values between structures are quite small, ranging from 0.217Å to 1.202Å. These values indicate that the native proteins and QTY analogues have similar structures. The superimpositions are A) BILF1^Nat^ *vs* BILF1^QTY^, B) BILF1^CyroEM^ *vs* BILF1^QTY^, C) LMP1^Nat^ *vs* LMP1^QTY^ Monomer, D) LMP1^CryoEM^ *vs* LMP1^QTY^ Dimer, E) LMP1^Nat^ *vs* LMP1^QTY^ Dimer, F) LMP2^Nat^ *vs* LMP2^QTY^.

Overall, the superposed structures exhibited high conformational similarity, as indicated by low RMSD values ranging from 0.217Å to 1.202Å. Notably, all superposed pairs except for BILF1^Nat^ vs BILF1^QTY^ (RMSD: 1.202Å) achieved RMSD values lower than 0.500Å. However, the relatively high RMSD of 1.202Å between structures of BILF1^Nat^ and BILF1^QTY^ does not undermine the effectiveness of the QTY code, because the superposition of BILF1^QTY^ with its Cryo-EM-determined native structure - BILF1^CryoEM^ - returned a significantly lower RMSD of 0.351Å. This indicates that the high RMSD of 1.202Å is likely due to lower confidence or prediction uncertainty in AlphaFold 3 predictions, rather than to true structural differences. Ultimately, these low RMSD values indicate that the water-soluble QTY analogs share remarkably similar structures with their native counterparts. Furthermore, these alignments provide additional evidence supporting the efficacy of the QTY code in preserving overall protein structure.

### Superpositions of experimentally determined structures of EBV integral membrane proteins, their computationally predicted native structures, and their water-soluble QTY analogs

For EBV integral membrane proteins with available experimental data, we performed three-way superpositions to compare further **i**) the experimentally determined EBV integral membrane protein structures, **ii**) the computationally predicted native structures, and **iii**) the computationally predicted water-soluble QTY analogs (Figure 4). Their excellent superpositions further confirm the notable structural similarities between native structures and their QTY analogs.

**Figure 4.**
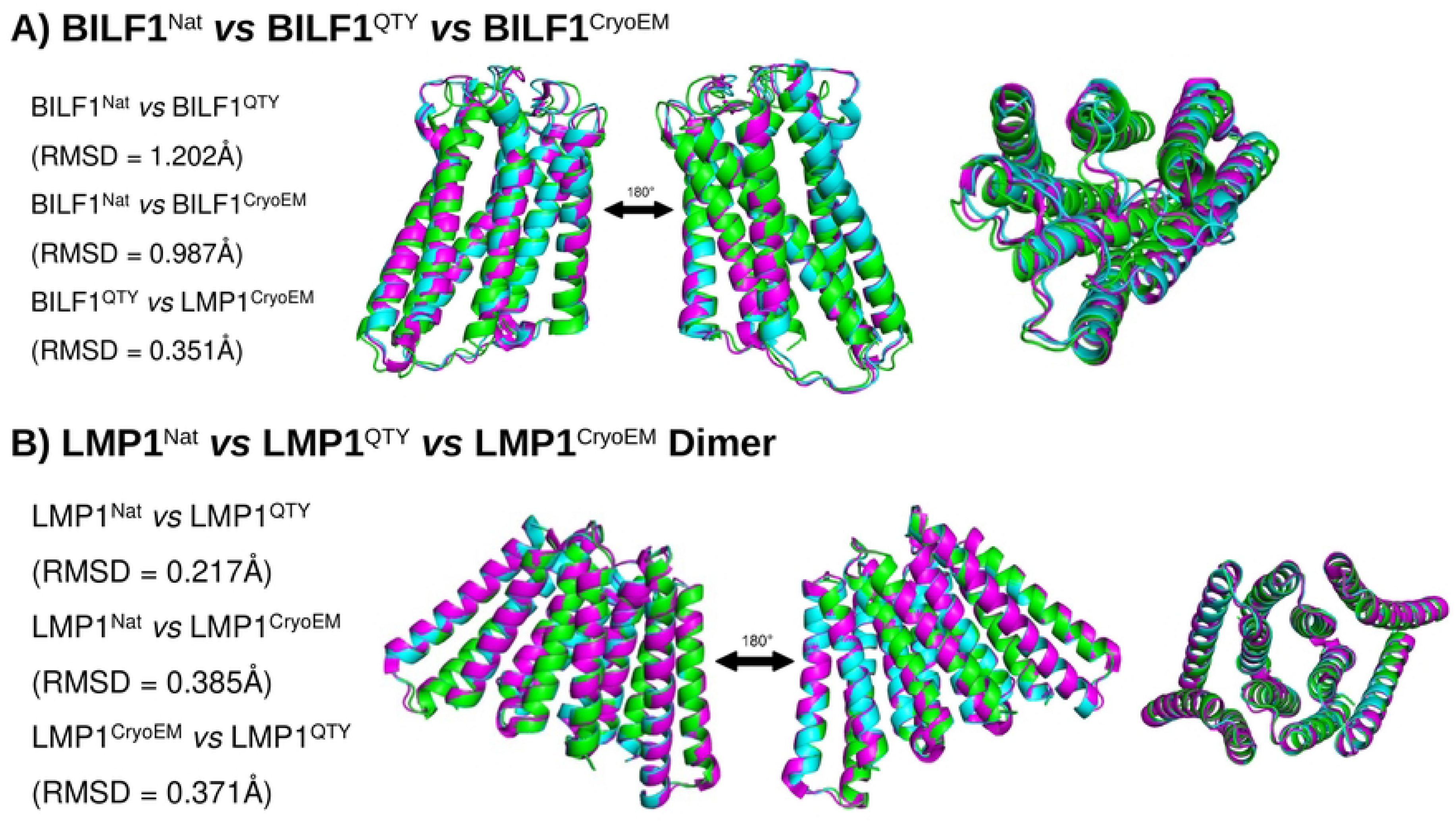
Three-way structural superimpositions of computationally predicted native (*green*), QTY-analogs (*cyan*), and experimentally determined Cryo-EM (*magenta*) models for two EBV integral membrane proteins. The three types of structures are superimposed very well, with largely insignificant differences between models, as supported by the low RMSD values, ranging from 0.217Å to 1.202Å. Three-way superimpositions of LMP2 and monomeric LMP1 are not displayed as they lack experimentally determined structures at the time of writing. The superimpositions are A) BILF1^Nat^ *vs* BILF1^QTY^ *vs* BILF1^CryoEM^, B) LMP1^Nat^ *vs* LMP1^QTY^ *vs* LMP1^CryoEM^ Dimer.

Furthermore, alignments between the predicted and experimentally obtained native structures yielded RMSD values below 1.000Å (BILF1^Nat^ vs BILF1^CryoEM^: 0.987Å, LMP1^Nat^ dimer vs LMP1^CryoEM^ dimer: 0.385Å). These minimal deviations corroborate the high predictive accuracy of AlphaFold3 and ColabFold for these specific viral proteins, thereby validating their use for protein modeling in this study.

### Analysis of the hydrophobic surface of native EBV integral membrane proteins and their water-soluble QTY analogs

Given the hydrophobic core of the phospholipid bilayer, TM domains of integral membrane proteins are typically highly hydrophobic [21]. EBV integral membrane proteins are no exception, containing hydrophobic TM alpha-helices consisting of large amounts of hydrophobic amino acids - leucine (L), isoleucine (I), valine (V), phenylalanine (F), methionine (M), tryptophan (W), and alanine (A). Since membrane proteins containing large numbers of these amino acids tend to expel water via lipid interactions, they readily aggregate and precipitate in aqueous environments. Consequently, detergents are required to solubilize these proteins, preserving their native structures and functions during isolation and purification [21].

The QTY code is presented as an engineering solution for designing detergent-free analogs of integral membrane proteins while preserving their structures and functions [23]. In Figure 5, hydrophobic regions of the protein surface are represented in yellow, while hydrophilic regions are shown in cyan. Since all six sets of compared structures show large patches of hydrophobic residues in the native protein structure being replaced by hydrophilic residues, we can reasonably conclude that QTY analogs have significantly increased water solubilities compared to their native counterparts. More importantly, the aforementioned superpositions demonstrate that the QTY analogs of EBV integral membrane proteins retain their tertiary structures despite the amino acid substitutions. These findings corroborate the efficacy of the QTY code and are consistent with our previous studies, including one on chemokine receptors, which demonstrated that QTY analogs retained ligand-binding activities, thermostabilities, and overall structures while becoming hydrophilic [23], [39].

**Figure 5.**
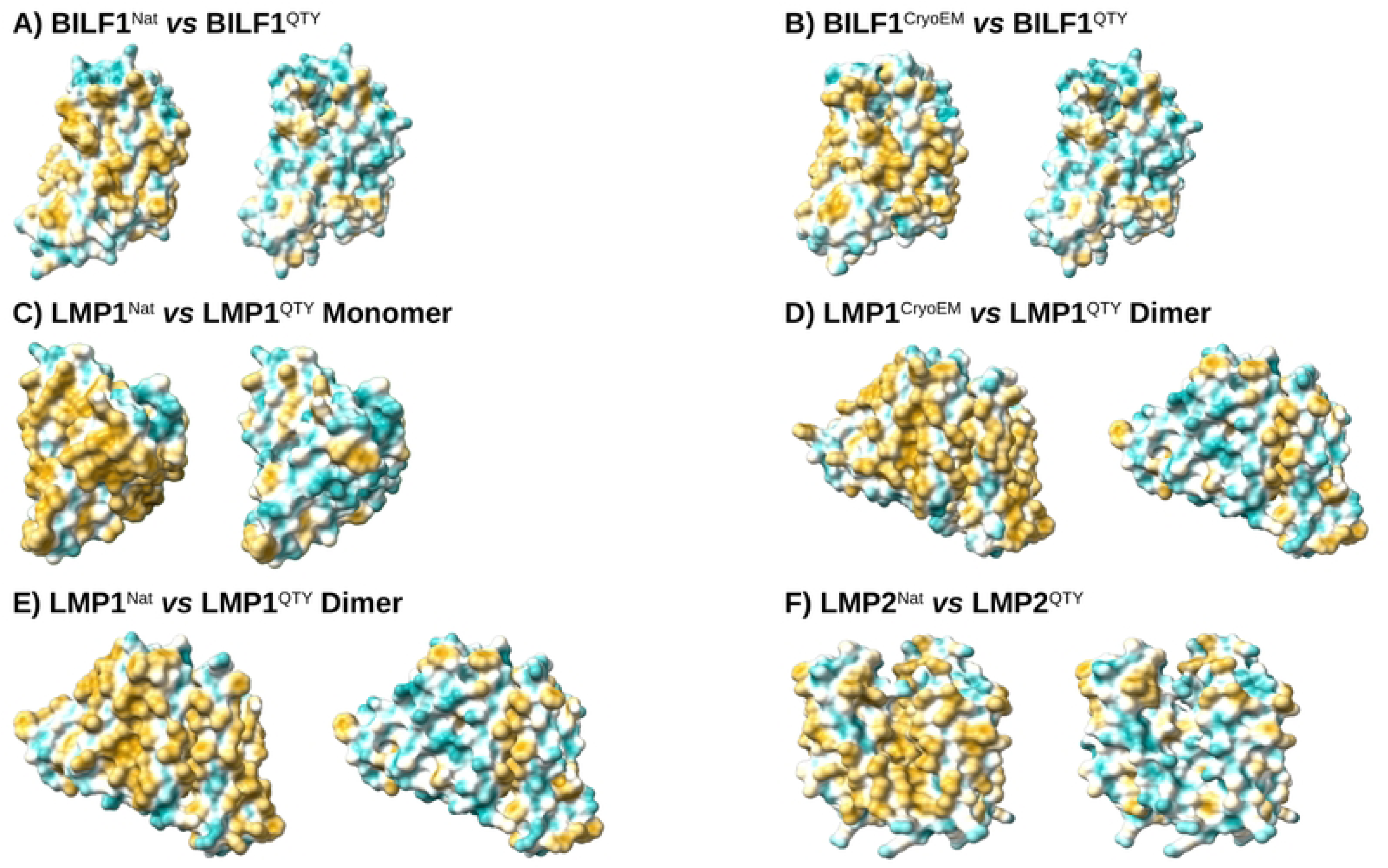
Hydrophobic surfaces of three native EBV integral membrane proteins (*left*) and their QTY analogs with reduced hydrophobicity (*right*). In the visualization above, hydrophobic regions are coloured yellow, while hydrophilic regions are coloured cyan. The native EBV integral membrane proteins have hydrophobic residues L, I, V, and F in their TM regions; the QTY code substitutes L, I/V, and F with the hydrophilic residues Q, T, and Y, respectively. After substitution, the hydrophobic TM domains become more hydrophilic with minimal changes to the proteins’ surface shapes. The disordered C- and N-termini were trimmed to improve structural clarity. The comparisons are A) BILF1^Nat^ *vs* BILF1^QTY^, B) BILF1^CyroEM^ *vs* BILF1^QTY^, C) LMP1^Nat^ *vs* LMP1^QTY^ Monomer, D) LMP1^CryoEM^ *vs* LMP1^QTY^ Dimer, E) LMP1^Nat^ *vs* LMP1^QTY^ Dimer, F) LMP2^Nat^ *vs* LMP2^QTY^.

To quantify the changes in hydrophobicity, CamSol intrinsic solubility profiles were generated for each EBV integral membrane protein and its respective QTY analog (Figure 6) [40]. All four analyzed structures show remarkable reductions in aggregation-prone regions, as indicated by shorter, fewer red regions with negative solubility scores. Moreover, there are longer and more blue regions with positive solubility scores. This overall increase in CamSol intrinsic solubility scores across all four proteins indicates reduced hydrophobicity and aggregation propensity of QTY analogs, highlighting their ability to remain more stable than their native counterparts in aqueous environments [41].

**Figure 6.**
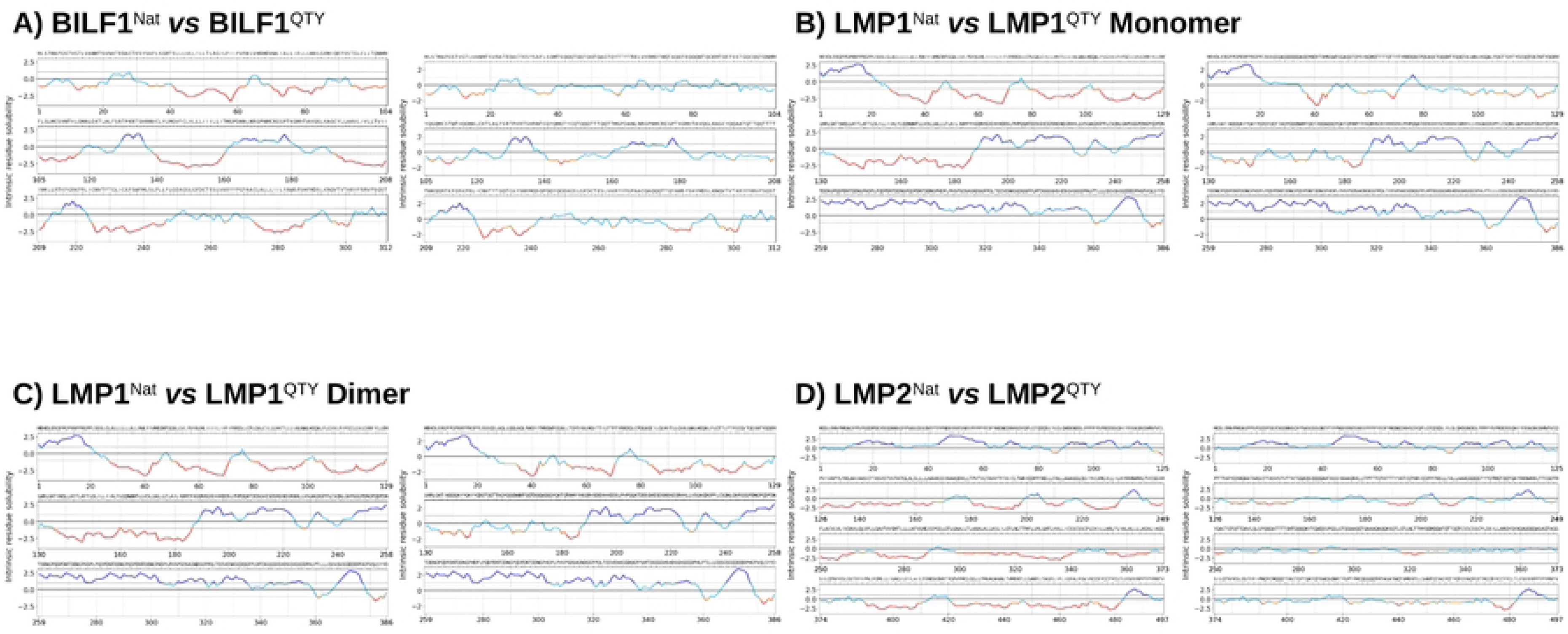
The CamSol Intrinsic solubility profiles of four native EBV integral membrane protein sequences (*Upper*) and their QTY analogs (*Lower*). In these profiles, insoluble and aggregation-prone segments are depicted in red, whereas highly soluble and aggregation-resistant regions are shown in blue. Across all four sequence pairs, applying the QTY code resulted in fewer red and more blue areas, demonstrating an overall reduction in hydrophobicity.

### Future directions and potential applications

Our study provides computational evidence that QTY-designed EBV integral membrane proteins are predicted to be water-soluble whilst retaining their original structures. If experimentally validated, these detergent-free BILF1, LMP1, LMP2 QTY analogs could provide useful tools for proteomics and pharmaceutical research.

First, by reducing the need for a detergent during protein purification and characterization, they may facilitate structural and functional studies of EBV integral membrane proteins. Allowing researchers to bypass the time-consuming step of detergent design and optimization would make research on BILF1, LMP1 and LMP2 more accessible, thereby facilitating future studies on EBV’s mechanisms of viral latency, oncogenesis, and immune evasion [22]. This is especially important for these proteins, as despite EBV’s widespread prevalence and their vital roles in EBV latency and pathogenicity, many of their functions remain unclear.

Second, as soluble analogs that preserve their native structures, QTY-designed EBV integral membrane proteins could accelerate structure-based drug design. Specifically, water-soluble QTY analogs could facilitate ligand-binding studies, epitope mapping, and screening assays for antibodies and inhibitors against EBV-associated therapeutic targets [42]. For instance, water-soluble QTY analogs of LMP1 may help identify molecules that interfere with its oligomerization, or its recruitment of transducer proteins TRAF and TRADD [11], [12]. Such applications could potentially lead to the development of new therapeutic treatment options against EBV.

Our validation of QTY analogs of BILF1, LMP1 and LMP2 remains at the structural bioinformatic level; therefore, future work should focus on experimental structural validation. Future studies could express and purify QTY analogs *in vitro*, then directly measure aqueous solubility and analyze their structures using Cryo-EM or similar techniques. Additionally, functional assays could be conducted to determine whether these QTY analogs retain key interactions with ligands.

## Conclusion

In our study, we used computational methods to design, predict, and validate water-soluble QTY analogs of BILF1, LMP1, and LMP2, three integral membrane proteins that play vital roles in the pathogenesis and latency of the Epstein-Barr virus. We first applied the QTY code to convert hydrophobic transmembrane alpha-helices into hydrophilic ones through systematic amino acid substitution [43]. Then, we compared molecular weights, isoelectric points, and intrinsic solubility scores, thereby providing computational evidence that the proteins exhibited significantly decreased hydrophobicity and aggregation propensity following QTY-design, with only minor changes in physical and chemical properties. The following RMSD analysis further suggested high structural similarity between the native and QTY-designed structures. Together, these findings corroborate the QTY code’s robust ability to design water-soluble analogs of integral membrane proteins, outlining a useful simple strategy for overcoming the dependence on detergents that currently limits accessibility of related research. Nevertheless, this study remains limited by our reliance on computational methods. Future work should therefore focus on experimental validation, including the expression and purification of these QTY analogs, and determining whether they maintain their interactions with ligands, other proteins, and signaling molecules.

## Methods

### Application of the QTY code, sequence alignments, and comparisons of other characteristics

The native amino acid sequences for the EBV integral membrane proteins BILF1, LMP1, and LMP2 were retrieved from the UniProt database (https://www.uniprot.org/) [44]. The transmembrane domain (TMD) sequences for each protein were determined through Protter 2D visualizations (https://protter.ethz.ch/start/) [45]. The QTY code was then applied to these TMD sequences using the Protein Solubilizing Server (https://pss.sjtu.edu.cn/) [46]. The server also calculated the molecular weights (MW), isoelectric points (pI), TMD variations, and overall variations of the native proteins and their respective QTY analogs. Protein sequence alignments were also generated using the Protein Solubilizing Server.

Moreover, the PDBePISA tool (https://www.ebi.ac.uk/pdbe/pisa/) was used to identify interfacing residues and residues making hydrogen/disulfide bonds, salt bridges, or covalent links (HSDC) when applying the QTY code to the LMP1 dimer [47]. These residues take crucial roles in maintaining the structural stability of the dimer, and therefore remain unchanged.

### In silico Predictions of Protein Structures

Due to the difficulty of modeling viral proteins *in silico*, multiple machine learning models were used to obtain the most accurate structures for each protein. Before in silico modeling, the highly disordered C- and N-termini of all three integral membrane proteins were truncated to further improve prediction reliability. The predicted Local Distance Difference Test (pLDDT) score was used as the primary metric to assess model confidence, with the highest-scoring model for each protein selected for further structural comparisons and analysis.

Specifically, *de novo* predictions were performed using AlphaFold3 (https://alphafoldserver.com/) for the protein structures BILF1^Nat^, BILF1^QTY^, LMP2^Nat^, and Boltz-2 (https://build.nvidia.com/mit/boltz2) for the LMP1^Nat^ monomer [34], [38]. Template-based modeling was utilized using ColabFold v1.6.1 (https://colab.research.google.com/github/sokrypton/ColabFold/blob/main/AlphaFold2.ipynb) to predict the LMP1^QTY^ monomer (template: LMP1^Nat^ monomer), LMP1^Nat^ dimer (template: 8XH6 from the RCSB PDB), LMP1^QTY^ dimer (template: 8XH6 from the RCSB PDB), and LMP2^QTY^ (template: LMP2^Nat^) [37].

### Structural Superpositions

Experimentally determined native structures of the EBV integral membrane proteins were retrieved from the RCSB PDB (https://www.rcsb.org/) when available [48]. The PDB files retrieved were: BILF1 (PDB ID: 7JHJ) and LMP1 dimer (PDB ID: 8XH6). The *in silico* predicted and available experimentally determined native structures were then superposed with their respective QTY analogs using PyMOL (https://www.pymol.org/), and the computed root mean square deviation (RMSD) values were recorded for further analysis.

### Structural Visualization

All structural superpositions and RMSD computations in this study were completed using PyMOL (https://www.pymol.org/) [49]. UCSF ChimeraX (https://www.cgl.ucsf.edu/chimerax/) was used to visualize surface hydrophobicity of the native protein structures and their QTY analogs [50].

### Hydrophobicity Analysis

To quantify the change in protein hydrophobicity after QTY design, the CamSol method was utilized through the CamSol Intrinsic Server (https://www-cohsoftware.ch.cam.ac.uk/index.php/camsolintrinsic) [40]. The method assigns each residue an intrinsic solubility score, generating a profile in which scores higher than 1 denote highly soluble regions, while scores lower than −1 denote poorly soluble, aggregation-prone regions. Besides, an overall intrinsic solubility score is computed for each protein, with lower scores indicating greater hydrophobicity [41].

